# Deep learning identifies pathological abnormalities predictive of graft loss in kidney transplant biopsies

**DOI:** 10.1101/2021.04.18.440166

**Authors:** Zhengzi Yi, Fadi Salem, Madhav C Menon, Karen Keung, Caixia Xi, Sebastian Hultin, M. Rizwan Haroon Al Rasheed, Li Li, Fei Su, Zeguo Sun, Chengguo Wei, Weiqing Huang, Samuel Fredericks, Qisheng Lin, Khadija Banu, Germaine Wong, Natasha M. Rogers, Samira Farouk, Paolo Cravedi, Meena Shingde, R. Neal Smith, Ivy A. Rosales, Philip J. O’Connell, Robert B. Colvin, Barbara Murphy, Weijia Zhang

**Author notes:** ZY and FS are co-first authors and contributed equally. **Correspondence:** Dr. Weijia Zhang, Division of Nephrology, Department of Medicine, Icahn School of Medicine at Mount Sinai, One Gustave L Levy Place, Box 1243, New York, NY 10029, Dr. Barbara Murphy, M.D. Division of Nephrology, Department of Medicine, Icahn School of Medicine at Mount Sinai, One Gustave L Levy Place, Box 1243, New York, NY 10029.

## Abstract

**Background:** Interstitial fibrosis, tubular atrophy, and inflammation are major contributors to renal allograft failure. Here we seek an objective, quantitative pathological assessment of these lesions to improve predictive utility.

**Methods:** We constructed a deep-learning-based pipeline recognizing normal vs. abnormal kidney tissue compartments and mononuclear leukocyte (MNL) infiltrates from Periodic acid-Schiff (PAS) stained slides of transplant biopsies (training: n=60, testing: n=33) that quantified pathological lesions specific for interstitium, tubules and MNL infiltration. The pipeline was applied to 789 whole slide images (WSI) from baseline (n=478, pre-implantation) and 12-month post-transplant (n=311) protocol biopsies in two independent cohorts (GoCAR: 404 patients, AUSCAD: 212 patients) of transplant recipients to correlate composite lesion features with graft loss.

**Results:** Our model accurately recognized kidney tissue compartments and MNLs. The digital features significantly correlated with Banff scores, but were more sensitive to subtle pathological changes below the thresholds in Banff scores. The Interstitial and Tubular Abnormality Score (ITAS) in baseline samples was highly predictive of 1-year graft loss (*p*=2.8e-05), while a Composite Damage Score (CDS) in 12-month post-transplant protocol biopsies predicted later graft loss (*p*=7.3e-05). ITAS and CDS outperformed Banff scores or clinical predictors with superior graft loss prediction accuracy. High/intermediate risk groups stratified by ITAS or CDS also demonstrated significantly higher incidence of eGFR decline and subsequent graft damage.

**Conclusions:** This deep-learning approach accurately detected and quantified pathological lesions from baseline or post-transplant biopsies, and demonstrated superior ability for prediction of posttransplant graft loss with potential application as a prevention, risk stratification or monitoring tool.

## Introduction

Kidney transplantation is the treatment of choice for patients with end-stage renal disease (ESRD)[1]. Interstitial fibrosis and tubular atrophy (IFTA) and inflammation are considered major contributors to post-transplant kidney allograft failure irrespective of etiology of injury[2]. Currently IFTA and inflammation are graded by pathological assessment of biopsies. While cumulative injury represented as categorical Banff scores have been associated with posttransplant graft function/survival, these have intermediate sensitivity for graft failure prediction in any given biopsy, due to inter- and intra-observer variability [3]. Prediction of long-term graft survival remains a major challenge. Post-transplant factors, such as the rate of decline of eGFR [4, 5] have shown predictive ability, however in the early transplant period, factors that predict the post-transplant course are lacking, yet are necessary to identify patients at risk of premature graft loss and guide subsequent patient management.

Recently, deep-learning-based approaches have been successfully applied to radiological medical images [6, 7] and histologically-stained images[8, 9], and studies in renal digital pathology have shown promise in detecting glomerular or interstitial abnormalities[10–15]. Good prediction of kidney tissue compartments [16–18] was obtained with pixel-level prediction algorithm U-Net [19]. An instance-level object detection algorithm mask R-CNN[20] was developed recently with advantages of performing object localization, shape prediction and object classification at the same time, which accurately distinguishes sclerotic from non-sclerotic glomeruli [21]. We reasoned that these deep-learning-based approaches could be applied for observer-independent histopathological assessment of transplant biopsies, offering distinct advantages for graft prognostication.

In this study, we first trained a deep-learning model based on both U-Net and mask R-CNN algorithms to accurately recognize normal/abnormal kidney tissue compartments and infiltrated MN leukocytes from both baseline (pre-implantation) and post-transplant biopsies. We then extracted slide-wide features to ensure capture of abnormalities in the interstitium, tubules, and inflammation and investigated their association with Banff scores and post-transplant graft outcomes in two large independent cohorts.

## Materials and Methods

### Study cohorts and biopsy slides

The Genomics of Chronic Allograft Rejection (GoCAR) [22] study is a prospective, multicenter study with patients have been followed for a median 5 years. AUSCAD is an Australia transplant cohort from Westmead Hospital, The University of Sydney NSW with patients being followed for a median duration of 4.5 years. In GoCAR, two protocol biopsy cores were taken from baseline (pre-implantation) or various times (3, 12, and 24m) post-transplant. One formalin-fixed, paraffin-embedded core was processed for histologic stains and scored centrally by at least 2 pathologists at Massachusetts General Hospital (MGH) according to Revised Banff 2007 Classification for Renal Allograft Pathology [23]. AUSCAD biopsy samples were formalin-fixed and paraffin-embedded prior to routine histological staining including Periodic acid–Schiff (PAS). All biopsies were scored locally according to the revised Banff 2007 classification for renal allograft pathology. Several of the AUSCAD biopsies were scored by both the MGH and Westmead pathologists to ensure there was consistency in diagnosis between the two centers. GoCAR slides were scanned with Aperio CS scanner at 20x objective with a 2x magnifier, AUSCAD slides were scanned by scanner from Hamamatsu company with a 20x objective.

PAS-stained slides in both cohorts were used in this study (Figure 1). Firstly, 93 slides that represented the spectrum of histological lesions were selected from 1164 PAS slides of biopsies taken from various time-points in entire GoCAR cohort. Sections of these slides covering glomeruli, interstitium, tubules, arteries and MNL infiltration were annotated under the guidance of pathologists to build deep learning detection models (training set, n=60; testing set, n=33). Secondly, the established pipeline was applied to 478 baseline whole slides (GoCAR, n=317; AUSCAD, n=161) and 311 12m post-transplant whole slides (GoCAR, n=200; AUSCAD, n=111) to extract digital features to be correlated with graft survival. These slides represented 404 patients from GoCAR cohort and 212 patients from AUSCAD cohort.

**Figure 1.**
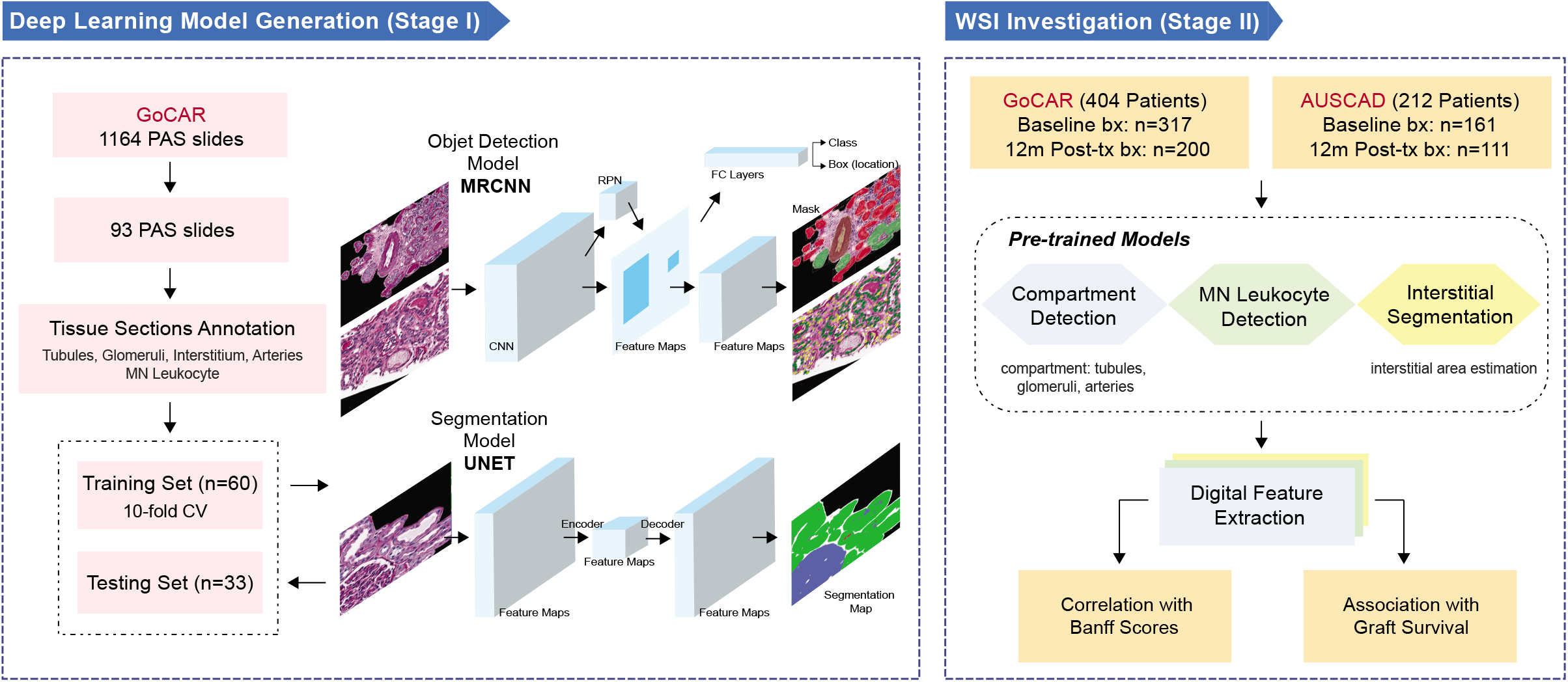
Study design. This study consists of two major stages. i) deep learning model generation. 93 slides that represented the spectrum of histological lesions were selected from GoCAR PAS slides and then randomly divided into discovery set (n=60) and testing set (n=33). The annotated sections of these slides were used for model construction and evaluation. During training process, we built the models based on two types of deep learning structures for compartment or MNL detection (by mask-RCNN) and tissue segmentation (by U-net). Models were determined through evaluation with 10-fold cross-validation and finally applied to the testing set. ii) WSI investigation. Using established deep learning model, we processed 789 baseline and 12m post-transplant WSIs from two independent cohorts (GoCAR and AUSCAD) and extracted a series of slide-wide digital features capturing the abnormalities in interstitium and tubules, and MNL infiltration. These features were further examined through association with Banff scores and post-transplant graft survival.

### WSI Deep learning analysis

The WSI (whole slide images) deep learning analysis procedure was divided into two stages: (I) deep-leaning-based detection model generation and (II) slide-wide feature extraction (the details were depicted in Figure 1, Figure S1 and described in Supplemental Method). Briefly, at Stage I, annotated PAS sections were pre-processed into 22,588 fixed-sized tiles through data augmentation. The deep learning model was generated and tuned on the training set (n=60 slides) with 10-fold cross-validation and the established model was applied to independent testing set (n=33 slides) for unbiased model evaluation. We constructed a compartment detection model and a MNL detection model using Mask R-CNN [20] and an interstitium estimation model using U-Net[19]. The detection accuracies were measured by True Positive Rate (TPR), Positive Predictive Value (PPV) and general *F_β_* score[24] where *β*=2. At Stage II, through scanning of unit window across the entire slide to identify interstitial/inflammatory Regions of Interest (ROIs), we defined slide-wide digital features capturing abnormalities in interstitium, tubules and MNL infiltration, which were then summarized into composite features reflecting overall kidney damages.

### Statistical analysis

Association of digital features with Banff scores/eGFR were measured by Spearman’s correlation. Association with graft loss were assessed by Cox proportional hazards regression. As for survival confounders adjustment, living/deceased donor, HLA mismatch and induction type were selected as confounders according to significance levels from univariate analysis.

## Results

### Demographic and clinical characteristics of study cohorts

We applied artificial intelligence techniques to PAS stained slides of kidney donor biopsies taken at baseline (pre-implantation) or 12m post-transplant in 404 patients from a multi-center international cohort (GoCAR)[22] and 212 patients from an external Australian cohort (AUSCAD) (Figure 1). The two populations had similar gender distribution, age and cold ischemia time (CIT), but they differed in ethnicity and clinical management protocols (Table 1). GoCAR patients had more diverse ethnic backgrounds including African-American/Hispanic (25% vs. none in AUSCAD), whereas AUSCAD recorded more deceased donors (78.77% vs. 53.71% in GoCAR). All AUSCAD patients received induction therapy predominantly with lymphocyte non-depleting agents (93.87%), whilst among 78.22% of GoCAR recipients who received induction, lymphocyte depleting agents (Thymoglobulin or Campath-1) were used in 39.36% and non-depleting agents in 38.86%. Overall the AUSCAD cohort had a lower graft loss rate (4.72% vs. 12.13% in GoCAR) during slightly shorter follow-up period (median 4.5 years vs. 5 years in GoCAR).

**Table 1.** Demographic and clinical characteristics in two independent kidney transplant cohorts.

| Characteristics | GoCAR (n=404)<br>Median±SD (%) | AUSCAD (n=212)<br>Median±SD (%) | P-value<sup>1</sup> |
| --- | --- | --- | --- |
| Recipient Age | 49.38±13.52 | 48.44±12.11 | 0.381 |
| Recipient Gender |  |  | 0.282 |
| Female | 129(31.93) | 77(36.32) |  |
| Male | 275(68.07) | 135(63.68) |  |
| Recipient Race |  |  | 1.7e-19 |
| White / Caucasian | 261(64.6) | 177(83.49) |  |
| Asian | 24(5.94) | 27(12.74) |  |
| African American | 76(18.81) | 0(0) |  |
| Hispanic | 25(6.19) | 0(0) |  |
| Other | 18(4.46) | 8(3.77) |  |
| Dialysis |  |  | 6.0e-04 |
| No | 89(22.03) | 23(10.95) |  |
| Yes | 315(77.97) | 187(89.05) |  |
| Kidney Disease |  |  | 1.3e-05 |
| Diabetes Mellitus | 139(34.41) | 79(37.98) |  |
| Glomerulonephritis | 74(18.32) | 66(31.73) |  |
| Hypertension | 77(19.06) | 14(6.73) |  |
| Polycystic Kidney Disease | 41(10.15) | 20(9.62) |  |
| Other | 73(18.07) | 29(13.94) |  |
| Donor Age | 42.02±15.51 | 45.43±16.8 | 0.019 |
| Donor Gender |  |  | 0.609 |
| Female | 197(48.76) | 105(50.97) |  |
| Male | 207(51.24) | 101(49.03) |  |
| Deceased Donor |  |  | 6.5e-10 |
| No | 187(46.29) | 45(21.23) |  |
| Yes | 217(53.71) | 167(78.77) |  |
| CIT minutes | 530.65±494.21 | 501.06±245.1 | 0.324 |
| HLA mismatch |  |  | 0.010 |
| 0 | 46(11.39) | 12(6.19) |  |
| 1-2 | 55(13.61) | 42(21.65) |  |
| 3-4 | 150(37.13) | 58(29.9) |  |
| 5-6 | 153(37.87) | 82(42.27) |  |
| DGF |  |  | 9.0e-04 |
| No | 334(82.67) | 150(70.75) |  |
| Yes | 70(17.33) | 62(29.25) |  |
| Induction Type |  |  | 5.1e-46 |
| Lymphocyte non-Depletion | 157(38.86) | 199(93.87) |  |
| Lymphocyte Depletion | 159(39.36) | 13(6.13) |  |
| None | 88(21.78) | 0(0) |  |
| Follow up Days | 1776.98±660.2 | 1637.39±849.81 | 0.038 |
| Death Censored Graft Loss |  |  | 0.002 |
| No | 355(87.87) | 202(95.28) |  |
| Yes | 49(12.13) | 10(4.72) |  |
| All Cause Graft Loss |  |  | 0.001 |
| No | 307(75.99) | 185(87.26) |  |
| Yes | 97(24.01) | 27(12.74) |  |

1. P-values are calculated by Fisher's exact test (categorical variables) or Student's t-test (continuous variables).

### Deep-learning-based WSI investigation defined abnormality in interstitium/tubules and MNL infiltration

Our two-stage study first generated a deep learning model detecting tissue compartments and mononuclear leukocytes (MNL), and then defined slide-wide abnormality features to be correlated with Banff scores[23] and graft outcomes (Figure 1, Supplementary Method). In Stage I, three types of models based on two deep-learning architectures were built on 60 slides (training set) using 10-fold cross-validation. The models respectively identified tissue compartments (tubules, glomeruli, etc.) and MN leukocytes (mask R-CNN), and interstitial area (U-Net). The final model was applied to an independent testing set (33 slides), and accurately recognized 96% of glomeruli, 91% of tubules and differentiated normal/abnormal tubules at True Positive Rate (TPR) of 81% and 84% respectively. Lastly, 85% and 96% of interstitial area and area covered by arteries were correctly identified (Table S1).

In Stage II, we created a pipeline which detected a series of slide-wide digital features specifically capturing abnormalities within biopsies (Figure S1A-S1B, Supplementary Method). *For quantifying abnormalities in tubules and/or interstitium*, we defined: i) Abnormal Interstitial Area Percentage, proportion of total abnormal interstitium area over WSI; ii) standardized Abnormal Tubule Density; iii) Interstitial and Tubular Abnormality Score (ITAS), a composite score of i) and ii). *To quantify inflammation in biopsies* (i.e. MNL infiltration), we defined: iv) MNL-enriched Area Percentage, proportion of MNL infiltration area over WSI; v) standardized MNL Density; vi) MNL Infiltration Score (MLIS), a composite score of iv) and v). Lastly, a Composite Damage Score (CDS), integrating both ITAS and MLIS, was defined as the estimation of overall graft damage. Figure 2A demonstrates an example application of our pipeline to an abnormal case: original WSI (a), whole slide prediction (b) and the masks highlighting abnormal interstitium/tubule regions (c) or MNL infiltration regions (d) which agreed with assessment by pathologists.

**Figure 2.**
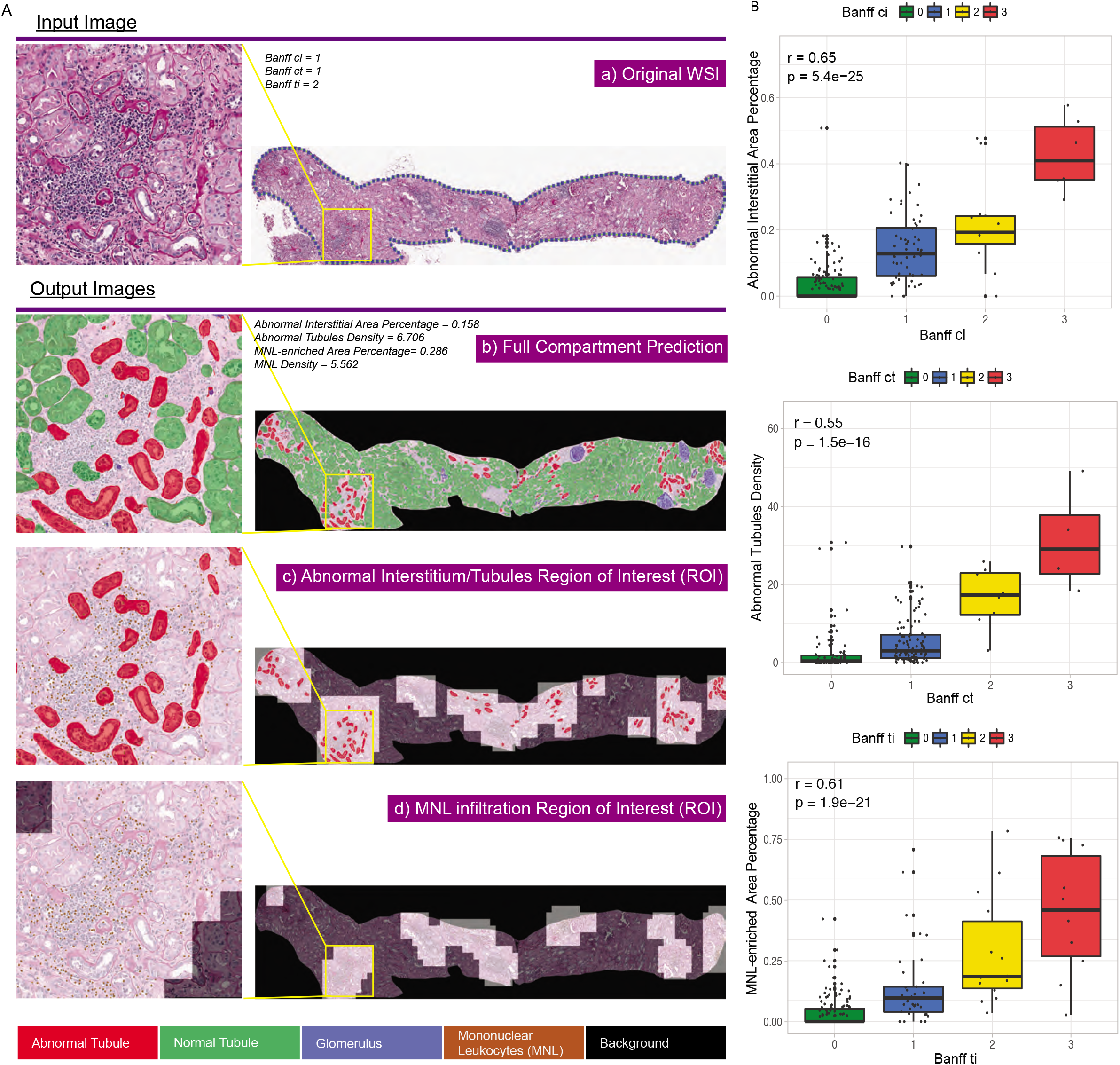
Demonstration of slide-wide digital features and correlation with corresponding Banff scores. **A)** Demonstration of slide-wide digital features from WSI investigation by an example WSI. a) original WSI; b) whole slide prediction; c) predicted abnormal interstitium/tubules regions of interest (ROI); d) predicted MNL infiltrated regions of interest (ROI). Left panel shows zoom-in inspections of one particular abnormal region within yellow box on WSI. **B)** Correlation of digital features with Banff scores. Correlation of Abnormal Interstitial Area Percentage and Banff ci score (top), Abnormal Tubules Density and Banff ct score (middle), MNL-enriched Area Percentage and Banff ti score (bottom) in GoCAR 12m post-transplant biopsy slides (n=200).

### Digital features were correlated with Banff scores

The Banff scores such as interstitial fibrosis (ci), tubular atrophy (ct) and total inflammation (ti) (graded by expert visual-assessment from different histological stains) are similar in pathological principle but different in quantification and technique to our PAS-based digital features (as illustrated in Figure S1B). Here, we examined the relationship between these two methods. We performed whole slide image (WSI) investigation extracting digital features in 789 WSIs from baseline (n=478) and 12m post-transplant (n=311) biopsies in both GoCAR and AUSCAD cohorts. Our data indicated that digital features (Abnormal Interstitial Area Percentage, Abnormal Tubules Density, and MNL-enriched Area Percentage) were significantly correlated with respective Banff scores in GoCAR baseline (Figure S2A) and 12m biopsies (Figure 2B). Similarly, the digital scores were correlated with Banff scores in AUSCAD (12m) where i+t was used because of unavailability of ti-score (Figure S2B).

Although highly correlated, we still identified discrepancies between the two scoring systems such as the case demonstrated in Figure S3A: here, Banff assessment reported all zeros but digital features indicated abnormal scores (illustrated by small clusters of shrunken tubules and MNLs). We then inspected all 137 cases classified as normal by Banff criteria (ci, ct, i, t, ti, g, cv=0) from baseline biopsies and identified 50 abnormal and 87 normal cases based on digital features. We discovered that the baseline digitally-abnormal group had significantly worse subsequent graft functions as measured by eGFR and significantly higher subsequent Chronic Allograft Damage Index (CADI) [25] scores post-transplant compared to normal group (Figure S3B). We also examined another set of 50 cases classified as normal by Banff criteria from 12m post-transplant biopsies and similarly identified 26 digitally-abnormal and 24 normal cases. No significant difference of subsequent graft outcomes was observed between 12m digitally-abnormal vs. - normal group, which could be due to limited case number. However, transcriptomic profiles of these patients in 3m post-transplant biopsies[22] revealed higher expression of immune response genes and lower expression of cell cycle, metabolic and transporter genes in 12m digitally-abnormal group, implicating ongoing interstitial/tubular injury or inflammation in histologically quiescent biopsies by Banff criteria (Figure S3C-S3D).

Taken together, the above data indicates that our digital features accurately reflect Banff scores and identified similar histological lesions. Furthermore, it suggested that in cases of discrepancy, digital quantitative scores offer a more sensitive assessment of graft damage below the Banff threshold.

### Baseline interstitial and tubular abnormality score predicted early graft damage and 1-year graft loss

The pathological evaluation of baseline biopsies could reveal donor kidney quality. However, its utility in post-transplant prognosis has been debated [26]. To explore a novel application for our digital features in baseline biopsies, we examined the association of individual/composite features with post-transplant graft failure and compared these with the performance of Banff-based scores. We also compared these with Kidney Donor Profile Index (KDPI), a composite demographic/clinical factor that is validated for deceased donors [27–29]. In GoCAR (n=317, Figure S4A, Table S2), we observed significant association of individual interstitial or tubular features, and composite ITAS with death-censored graft loss (DCGL) and all-cause graft loss (ACGL) in univariate or multivariate Cox models. In AUSCAD (n=161, Figure S5A, Table S3), the association with graft survival was confirmed in ACGL but not in DCGL, which could be due to fewer DCGL cases.

Time-dependent AUC estimation in GoCAR indicated that baseline individual or composite digital features outperformed individual Banff scores or ci+ct respectively in prediction of DCGL within 12m (Figure 3A). Next, we divided baseline biopsies into three risk groups by composite feature ITAS: high (ITAS>0.6), intermediate (0.1<ITAS ≤ 0.6) and low (ITAS ≤ 0.1) risk. The high/intermediate ITAS risk groups exhibited significantly higher DCGL rates compared to the low ITAS risk group over the entire period of follow up. These differences were most apparent in the first 12 months post-transplant (Figure 3B, *p*=2.8e-05) and in the deceased-donor sub-cohort (Figure S4B, *p*=5.8e-03). ITAS was superior to KDPI (Figure S4C, *p*=0.132, KDPI>85%, 20%<KDPI≤85%, KDPI≤20%) for risk stratification of DCGL. Of note, a significantly higher ACGL rate was also observed in baseline high ITAS risk group (Figure S4D), whilst high and intermediate ITAS risk groups demonstrated a sustained decline in eGFR over the first 12m posttransplant (Figure 3C), consistent with incrementally significant correlation of ITAS with posttransplant eGFR at 3m (*p*=0.001), 6m (*p*=7.6e-05), and 12m (*p*=1.5e-05)). A significantly higher incidence of delayed graft function (DGF) (*p*=3.9e-05), and early (3-month post-transplant) graft damage as measured by the CADI score>2 (*p*=0.002) were observed in high/intermediate ITAS risk groups (Figure 3D). In AUSCAD (n=161), the association of ITAS risk groups with graft loss and other clinical outcomes were validated as shown in Figure S5B-S5D.

**Figure 3.**
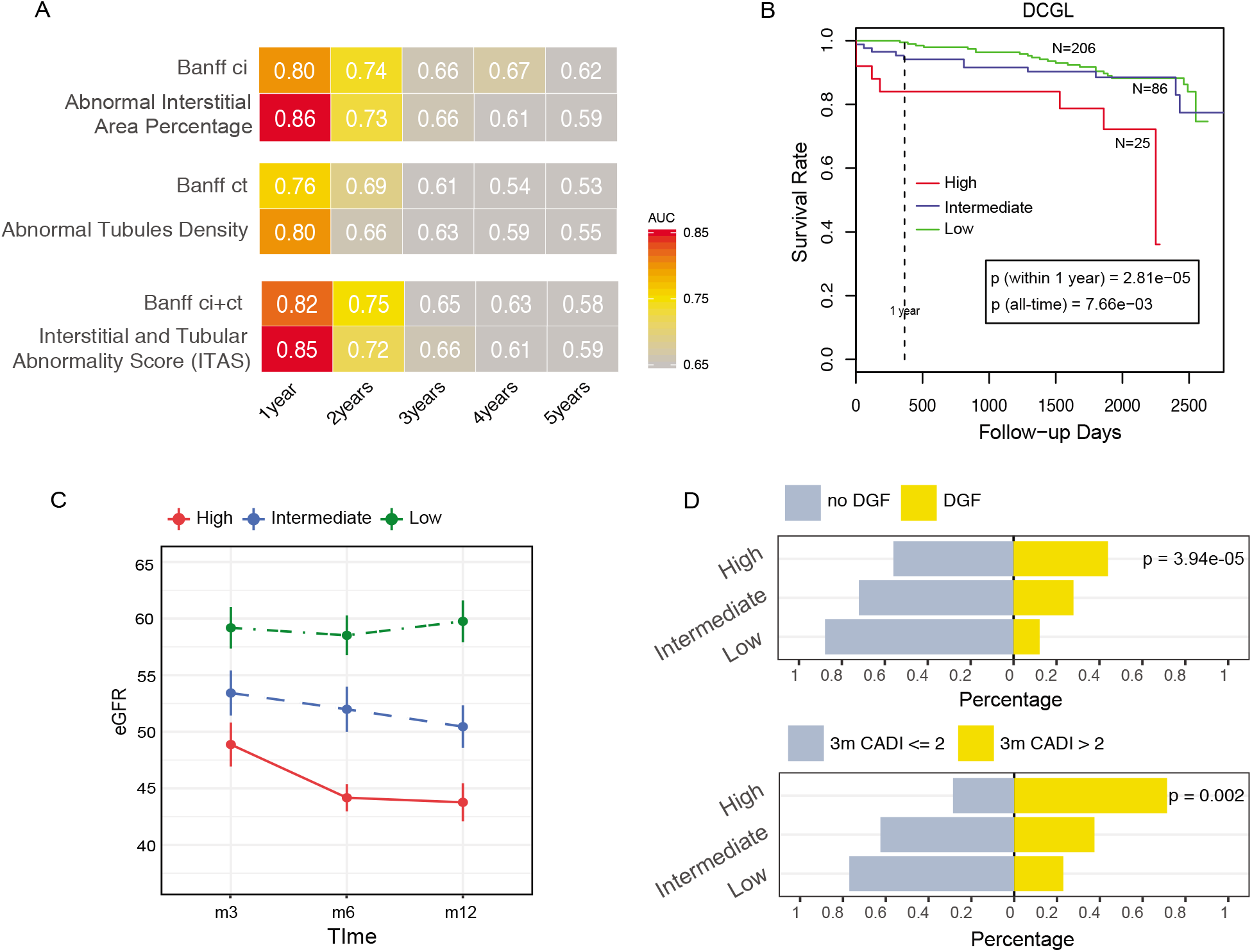
Association of baseline digital features with post-transplant graft outcomes in GoCAR cohort. **A)** Heatmap of time-dependent AUCs in predicting death-censored graft loss (DCGL) by Banff scores and digital features at different time intervals in baseline biopsy slides (n=317). Numbers and yellow-red color range of boxes represent AUC values at given time points. **B)** Kaplan-Meier curves of DCGL in high, intermediate and low risk groups stratified by Interstitial and Tubular Abnormality Score (ITAS) from baseline biopsies (n=317). Baseline ITAS groups are defined as: high: ITAS>0.6, intermediate: 0.1<ITAS≤0.6, low: ITAS≤0.1. P-values are calculated by log-rank test. **C)** Average eGFR values over time within 12m post-transplant per baseline ITAS risk group. Error bars represent 0.1x standard deviation from mean values. **D)** Bar charts demonstrating proportions of DGF/no DGF (upper) and 3m post-transplant CADI >2/≤2 (lower) among three baseline ITAS risk groups. P-values are calculated by Fisher’s exact test.

### 12m post-transplant composite damage score predicted long-term graft loss

Since our data showed that composite baseline digital score predicts early but not long-term DCGL, we examined longer term subsequent graft survival utilizing 12m post-transplant biopsy slides in both cohorts. In GoCAR (n=200, Figure S6A, Table S4), digital interstitial and tubular features, superior to corresponding Banff ci, ct scores, were significantly associated with long-term DCGL and ACGL with or without adjustment for clinical confounders (living/deceased donor, HLA mismatch, and induction type), while the MNL feature was comparable to the Banff ti score in association with DCGL. The associations of 12m digital features with long-term survival were validated in AUSCAD (n=111, Figure S7A, Table S5).

We observed that 12m digital features outperformed corresponding Banff scores including CADI in predicting long-term graft loss with superior time-dependent AUCs in GoCAR (Figure 4A). We then utilized the Composite Damage Score (CDS) summarizing abnormalities detected in interstitium, tubules, and inflammation for graft loss risk stratification. A 12m CDS >1.5 surpassed other clinical factors (>30% 3m to 12m eGFR decline, 3m or 12m acute cellular rejection) in longterm survival prediction (Figure 4B). Kaplan-Meier curves of DCGL (Figure 4C, *p*=7.3e-05) and ACGL (Figure S6B, *p*=1.6e-06) confirmed significantly lower survival rate in patients with high 12m CDS. We also identified significant associations of 12m CDS risk groups with other published surrogate outcomes including >30% 6m to 24m eGFR decline[4, 5] (*p*=0.010) and progressive histologic damage (*p*=0.005, 24m-CADI>2) (Figure 4D). These analyses in AUSCAD (n=111) also validated the predictive ability of 12m CDS for long-term survival (Figure S7B-S7E). Thus, high 12m CDS (>1.5), obtained at 12m post-transplant, is an alternative surrogate for long-term graft loss.

**Figure 4.**
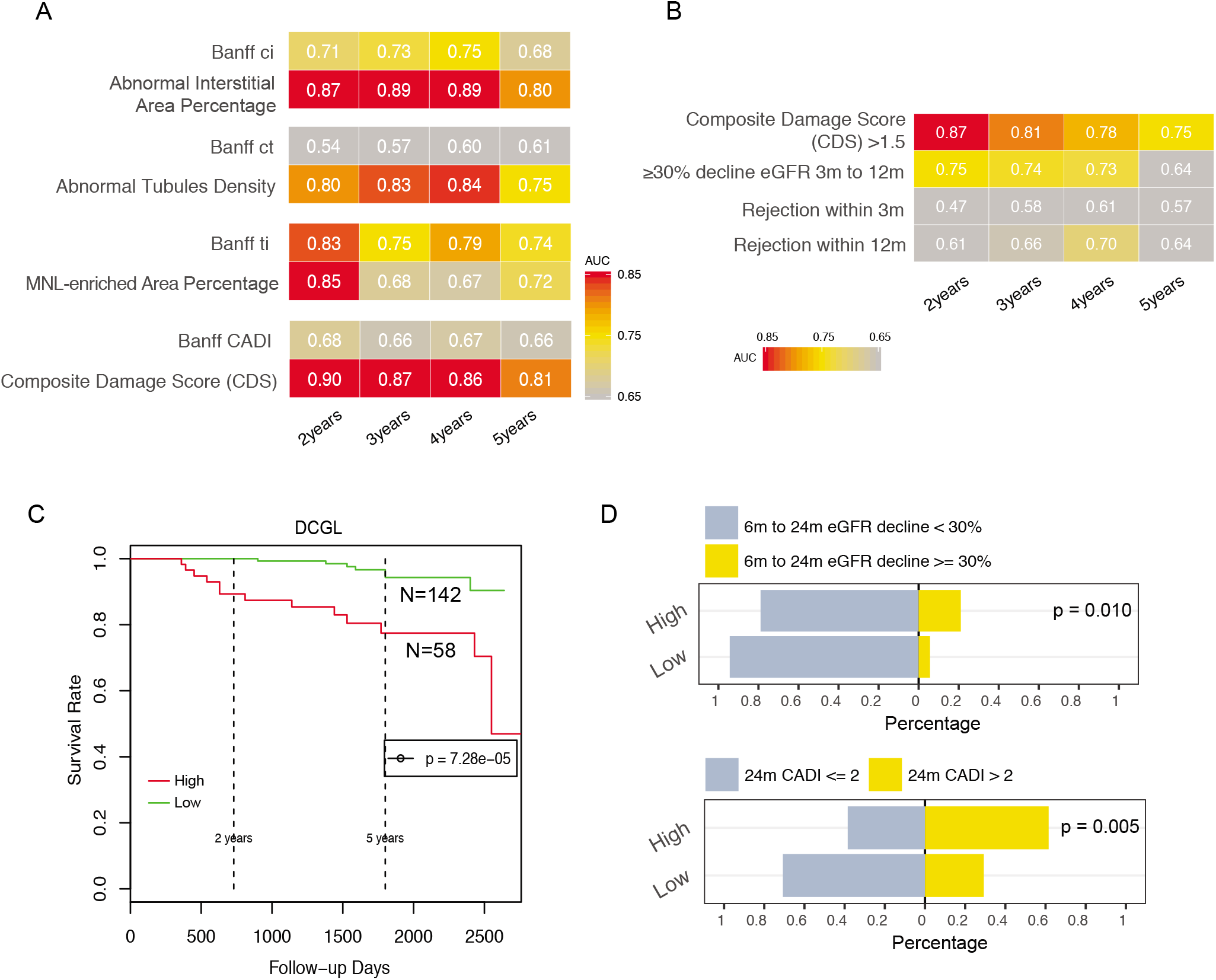
Association of 12m post-transplant digital features with post-transplant graft outcomes in GoCAR cohort. **A)** Heatmap of time-dependent AUCs in predicting death-censored graft loss (DCGL) by Banff scores and digital features at different time intervals in 12m posttransplant biopsy slides (n=200). Numbers and yellow-red color range of boxes represent AUC values at given time points. **B)** Heatmap of time-dependent AUCs in predicting DCGL by 12m Composite Damage Score (CDS, capturing the interstitial and tubular abnormality and MNL infiltration) high/low group and other clinical factors which were obtained prior to or at 12m. 12m CDS groups are defined as: high: CDS>1.5, low: CDS≤1.5. **C)** Kaplan-Meier curves of DCGL in high and low risk groups stratified by 12m CDS. P-value is calculated by log-rank test. **D)** Bar charts demonstrating proportions of 6m to 24m eGFR decline ≥30%/<30% (upper) and 24m posttransplant CADI >2/ ≤ 2 (lower) between 12m CDS risk groups. P-values are calculated by Fisher’s exact test.

## Discussion

We constructed a deep-learning-based histopathologic assessment model recognizing and quantifying interstitial, tubular, and inflammatory abnormalities in kidney transplant biopsies. WSI investigation of baseline and 12m post-transplant biopsies validated these digital features and further explored potential applications of composite features in clinical practice. Our digital features not only exhibited strong correlation with relevant Banff scores, but also detected subtle changes below the thresholds in Banff scores. Composite features of baseline ITAS and 12m CDS were identified to be predictive of early and late graft outcomes respectively, implying utility in transplant prognosis. To the best of our knowledge, this is the first study applying artificial intelligence techniques in identifying digital pathological features associated with solid organ transplant survival from both baseline and post-transplant biopsies with validation in multiple prospective cohorts.

Compared to previous investigations in deep-learning-based kidney tissue compartment detection [16–18], our study advances the field in four ways: i) Besides U-Net, we incorporated a mask R-CNN architecture for more efficient and accurate detection of the normal/abnormal compartments. ii) As inflammation is another major contributor to graft failure, we added a mask R-CNN based MN leukocyte detection model in post-transplant biopsy evaluations, improving graft loss predictive ability. iii) The slide-wide pathological lesions were quantified through definition of individual features in interstitium, tubules and MNL infiltration respectively, or composite features reflecting overall kidney damage. iv) We explored a novel clinical application of developed digital features for graft survival prediction in two well-designed cohorts. Both GoCAR and AUSCAD are large prospective cohorts which collected protocol biopsies pre-implantation and at time-points post-transplant, and followed up for median 4.5-5 years. Biopsies in GoCAR were graded centrally by at least 2 pathologists at MGH to minimize assessment variations.

Although many attempts have been made, no consistent association has been established between baseline histological findings and post-transplant outcomes among publications[26, 30]. Two limitations of these approaches are inter-operator variability due to inconsistency in histological assessment and variable expertise in transplant pathology[30, 31]. The Banff system itself has limitations by using categories rather than continuous variables. Our machine-based process overcomes these drawbacks by producing consistent and automated results in <30 minutes given the same scanning slide. The ITAS at baseline, was superior to Banff ci+ct and KDPI, and demonstrated the ability of stratifying risk of early graft damage, thus providing early information with utility for post-transplant monitoring, risk stratification or potential interventional trials.

We identified that the Composite Damage Score (CDS) from 12m protocol biopsies predicted long-term graft survival, outperforming histology and clinical factors. Reporting longer term hard outcomes from prospective trials has been an issue in kidney transplantation research[32]. The identification of surrogate endpoints is a major unmet need that often prevents the design of adequately powered trials. Recent studies proposed using eGFR decline within 24m/36m as a long-term graft loss surrogate[4, 5]. However, such a surrogate has the following limitations: i) Creatinine measurement is impacted by a number of factors including timing of collection in the day, diet, and inter laboratory variation. [33, 34]; ii) eGFR decline has low detection sensitivity because it requires multiple measurements during long-term follow-up, and the ≥40% decline from 6m to 24m, as suggested by a prior study for graft loss prediction [5], only occurred in 4% of patients in the GoCAR cohort although rates of graft loss were 12% for DCGL and 24% for ACGL. In contrast, 12m CDS was able to detect 29% of GoCAR and 21% of AUSCAD population as high risk as early as 12m while still exhibiting optimal AUCs in long-term graft loss prediction.

Our study has limitations. First, the identification of microvascular inflammation (g and ptc) and arteritis (v) requires further refinement. Further, current MN leukocyte detection score appears less accurate when detecting leukocytes within tubules than in interstitium. Its ability to diagnose and grade acute cellular rejection has not been demonstrated and it is unable to differentiate between antibody and cell mediated rejection. In addition, further refinements are required to diagnose transplant glomerulopathy or de novo or recurrent glomerular diseases. We aim to improve the model by integrating abnormality detection from other compartments and additional histological slides specifically with microvascular inflammation to extend the capture of pathological lesions.

In summary, our deep learning approach provided a reliable risk stratification of post-transplant graft survival using transplant biopsies at baseline and 12m post-transplant. This represents a novel and reproducible approach to facilitate early prevention, risk stratification or post-transplant monitoring in clinical practice.

## Supporting information

all

## Author Contributions

Z.Y. designed and performed computational analyses, and drafted the paper. F.Salem supervised pathology annotation and interpretation, and edited the paper. M.C.M contributed to study design and edited the paper, K.K. was involved clinical data collection of AUSCAD cohort and edited the paper; C.X., M.RHA.R, L.L., F.Su and Z.S. annotated slides; S.H. was involved clinical data and image collection of AUSCAD cohort; C.W., H.W., S.F., Q.L. K.B. and S.F. helped the clinical data mining and interpretation for GoCAR cohort, and critical reading of the manuscript; G.W. and N.M.R were involved in patient and sample management for AUSCAD cohort. P.C. was involved in data interpretation and edited the paper; M.S. was involved in AUSCAD cohort pathological assessment. R.N.S, I.A.R, were involved in pathological assessment of GoCAR cohort. P.O.C supervised ASUCAD cohort study and edited the paper. R.B.C supervised the pathology in GoCAR cohort and edited the paper; B.M. supervised GoCAR cohort study and was involved in study conception and paper editing. W.Z. conceptualized and designed this study and edited the paper.

## Data Sharing

The code and de-identified participant data will be made available to qualifying researchers by requesting to corresponding authors. Proposals will be reviewed by the investigators and collaborators based on scientific merit. If the proposal is approved, the data will be shared through a secure data transfer site.

## Acknowledgement

We thank Ms. Meyke Hermsen in Dr. Jeroen A. W. M. van der Laak’s lab in Department of Pathology of Radboud University Medical Center in Nijmegen for suggestions in kidney compartment annotation using ASAP (Automated Slide Analysis Platform) program. We thank Scientific Computing Division at the Icahn School of Medicine at Mount Sinai for providing computational resource.

