## Supplementary material for "Deep learning identifies pathological abnormalities predictive of graft loss in kidney transplant biopsies": all

### Supplementary Appendix

#### Supplementary Method

##### Image pre-processing

We annotated sections from 93 GoCAR[1] PAS stained slides for tissue compartment and monolayer leukocytes (MNL) prediction using ASAP software (https://computationalpathologygroup.github.io/ASAP/) under the guidance of pathologists. Each section was outlined by a boundary and the area within boundary but apart from annotated objects were defined as interstitium by default. The raw sections were divided into 22588 fixed-sized tiles at 20x objective-power and then transformed by data augmentation process including position shifting, rotating, flipping, perspective transforming, color transferring, contrast transforming, and noise feeding. Every fixed-sized tile was paired with a ground truth classification image labeling group information at the same size for model evaluation. These two types of images were served as input in model construction process.

##### Deep-learning model generation

We split 93 slides roughly in the ratio 2:1 into discovery set (n=60) and testing set (n=33). Discovery set were used for model construction in training process and testing set were used for final evaluation and were kept untouched during training process. We performed 10-fold cross-validation within discovery set thus divided the discovery set into 10 equal sized portions. During each model training process, we used 9/10 of samples as training set and the left-out 1/10 of samples as validation set to tune the model. As a result, 10 separate base models were created based on 10 partitions and the final prediction were made by aggregating results from each base model.

Two types of CNN structures were used in this study: a model segmenting images at instance level: Mask Region-based Convolutional Neural Network (Mask R-CNN)[2] and a model segmenting images at pixel level: U-Net[3]. The former model Mask R-CNN first extracts feature maps from input images by a convolutional backbone structure and generates region proposal of given objects through Region Proposal Network (RPN). These proposed regions are later passed through another neural network to generate multi-categorical classes, bounding boxes and masks for objects. The U-Net model on the other hand makes predictions for each pixel on input images instead of instance. It has a symmetrical "U" shape architecture consisting of an encoder which extracts features from input images by convolution blocks and a decoder which expends contracted vector back to segmentation map at input image size. The number of blocks in encoder step is as same as the number of blocks in decoder step.

We constructed a compartment detection model and a MN leukocyte detection model using Mask R-CNN structure[4] and a tissue segmentation model using U-net structure[5]. Specifically, the compartment detection model was trained for 90 epochs with batch size of 10 using 1024x1024 pixels input images; the MN leukocyte detection model was trained for 250 epochs with batch size of 6 using 512x512 pixels input images; the tissue segmentation model was trained for 100 epochs with batch size of 8 using 512x512 pixels input images. The detection model used RestNet-

101 as backbone and was tuned based on pre-trained weights from MS COCO dataset. We applied GSD optimizer with learning rate of 0.001 and momentum of 0.9. Total loss was calculated as the sum of loss of RPN classifier, RPN bounding box, MRCNN classifier, MRCNN bounding box, and MRCNN mask, where cross-entropy was used for classification problems and smooth L1-loss was used for bounding box refinement. The segmentation model was constructed using 3 down-sampling layers and 3 up-sampling layers and the first layer contained 32 feature maps. ADAM optimizer was chosen for weight updating at learning rate of 0.001 and cross-entropy was used for loss function. Best epoch/model was determined by evaluating loss from each training and validation partition. Finally, we applied the best model to testing data set for unbiased model evaluation. Accuracies were measured by True Positive Rate (TPR) and Positive Predictive Value (PPV) and general $F_\beta$ score[6] where $\beta$=2.

The GPU machine we used was equipped with 36 Intel(R) Xeon(R) W-2195 CPUs (18 cores), 128 GB Memory, and 4 GPUs of Quadro RTX 8000. All processes ran on Ubuntu 18.04 system.

**Whole slide investigation and digital feature definition**

We applied above three models to whole slide images (WSI) and assembled results into full prediction masks including all tissue compartment objects and MN leukocytes. Since this study focused on the features in cortex, medullar region and adjacent artery along with imperfectly cut or scanned fragments were excluded in WSI analysis. Due to the instance and pixel level prediction nature of MRCNN and U-Net, we were able to perform object counting as well as area estimation. In general, from WSI prediction, we defined basic features such as the size of a slide, the number of glomeruli and tubules, the percentage of glomeruli, tubules and interstitial area over slide, as well as a series of abnormal features focusing on interstitial space, abnormal tubules and inflammation.

To define slide-wide abnormal features, we first introduced the concept of Region of Interest (ROI) window to identify local abnormal regions with respect to interstitium and inflammation (Figure S1A). Given a whole slide prediction image, we applied a 384x384 pixels unit window sliding over the image with stride of 128 pixels. Within each unit window, we examined three statistics and defined two types of ROI: **interstitial ROI** and **inflammatory ROI**. A unit window was determined interstitial ROI if it had wide interstitial space but narrow space of background noise in tubule-enriched regions as defined as: $Sparsity(I) > 0.35$, and $Density(O) < 0.2$, and $Density(B) < 0.2$, where

$$Sparsity(I) = \frac{area(Interstitial\ Space)}{area(Interstitial\ \ Space)+area(Tubule)} \text{ per unit window} \quad (1)$$

$$Density(O) = \frac{area(Glo)+area(Other\ groups)}{384\times384} \text{ per unit window} \quad (2)$$

$$Density(\mathrm{B}) = \frac{\mathrm{area(backgroud)}}{384\times384} \text{ per unit window} \quad (3)$$

A unit window was determined inflammatory ROI if it had enriched mononuclear leukocytes (MNL) as $Density(MNL) > 43$, where

$$Density(MNL) = N(\mathrm{MNL}) \text{ per unit window} \quad (4)$$

After one slide is processed with sliding-window scanning, the pipeline generated two types of ROI masks highlighting abnormal interstitium area and MN leukocytes infiltration area, respectively.

Abnormal features were then defined in terms of interstitial space, abnormal tubules and MN leukocytes infiltration at WSI level or ROI level:

- Interstitium features: Feature (5) estimates overall percentage of intestinal space over WSI area.

$$\boldsymbol{Abnormal\ Interstitial\ Area\ Percentage} = \frac{area(Interstitial\ Space\ within\ \boldsymbol{interstitial\ ROI})}{area\ (WSI)} \quad (5)$$

- Abnormal tubules feature: Feature (6) summarizes number of abnormal tubules per 1000x1000 unit area.

$$\boldsymbol{Abnormal\ Tubules\ Density} = \frac{N(Abnormal\ Tubules)}{area(WSI)} \times 10^6 \quad (6)$$

- Inflammation features: Feature (7) estimates proportion of MN leukocyte enriched area over WSI area. Feature (8) summarize average MN leukocyte number per inflammatory ROI.

$$\boldsymbol{MNL\text{-}enriched\ Area\ Percentage} = \frac{area(\boldsymbol{Inflammatory\ ROI})}{area(WSI)} \quad (7)$$

$$\boldsymbol{MNL\ Density}\ (\boldsymbol{infR}) = Average\ N(MNL)\ weighted\ across\ \boldsymbol{Inflammatory\ ROIs} \quad (8)$$

Basically, the digital features were defined in consideration of two aspects: i) how widespread is given abnormal features over slide such as Abnormal Interstitial Area Percentage (5) and MNL-enriched Area Percentage (7); ii) how dense is given abnormal objects per unit area such as Abnormal Tubules Density (6) and MNL Density (infR) (8).

Moreover, we integrated above individual features into composite scores:

$$\boldsymbol{MNL\ Infiltration\ Score}\ (\boldsymbol{MLIS}) = \max\left(\boldsymbol{MNL\text{-}enriched\ Area\ Percentage} \times log_2\left(100\boldsymbol{MNL\ Density}\ (\boldsymbol{infR})\right),\ 0\right) \quad (9)$$

$$\boldsymbol{Interstitial\ and\ Tubular\ Abnormality\ Score}\ (\boldsymbol{ITAS}) = \max\left(\boldsymbol{Abnormal\ Interstitial\ Area\ Percentage} \times log_2(10\boldsymbol{Abnormal\ Tubules\ Density}),\ 0\right) \quad (10)$$

$$\boldsymbol{Composite\ Damage\ Score}\ (\boldsymbol{CDS}) = \boldsymbol{MLIS} + \boldsymbol{ITAS} \quad (11)$$

MLIS integrates both MN leukocyte coverage and density over WSI; ITAS reflects combined abnormality of interstitium and tubules over WSI; CDS represents overall graft damage with regard to interstitium, tubular abnormalities and inflammation over WSI.

In summary, our whole slide investigation pipeline generated three types of outputs for one WSI: i) whole slide prediction masks demonstrating kidney tissue compartments within a slide. ii) two ROI masks representing interstitium and inflammation abnormality. iii) a comprehensive data report summarizing individual or composite features.

**Microarray analysis**

Microarray data from GoCAR 3m post-transplant biopsies were described in our previous published paper[1]. Differential expression analysis was conducted by limma test[7] and significant genes were subjected to Gene Ontology [8] enrichment analysis by Fisher's exact test.

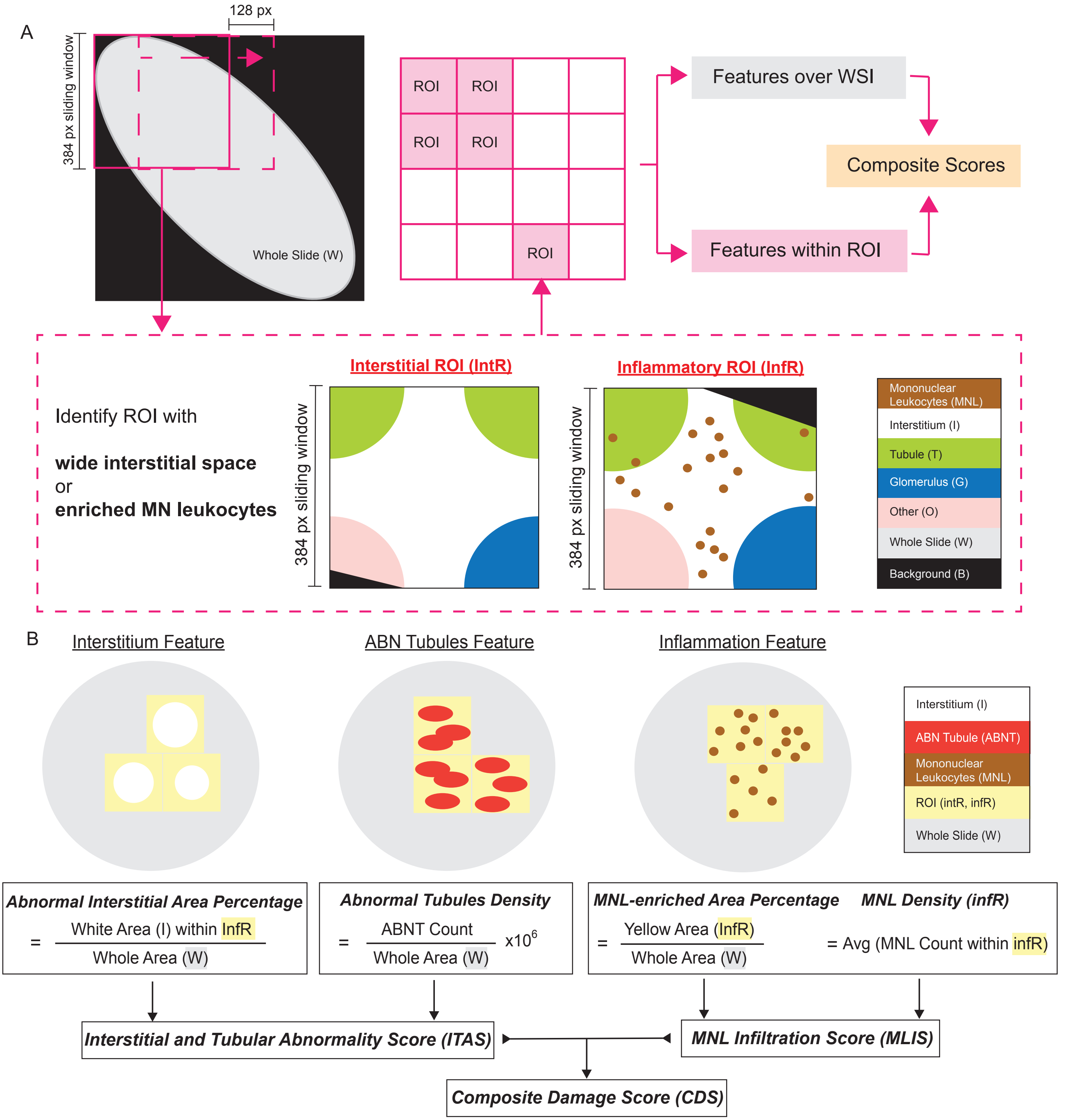

**Figure S1. Illustration of slide-wide digital features extraction process and definition. A)** Illustration of feature extraction process. We used a 384x384 pixel unit window scanning across WSI at stride of 128-pixel distance. Windows that had wide interstitial space or high amount of MN leukocyte were defined as interstitial regions of interest (intROI) or inflammatory regions of interest (infROI). A series of individual features were defined at ROI or slide level and further integrated into composite scores aiming for overall abnormality estimation. **B)** Illustration of definition (calculation) of individual digital features in interstitium, tubules and MNL infiltration.

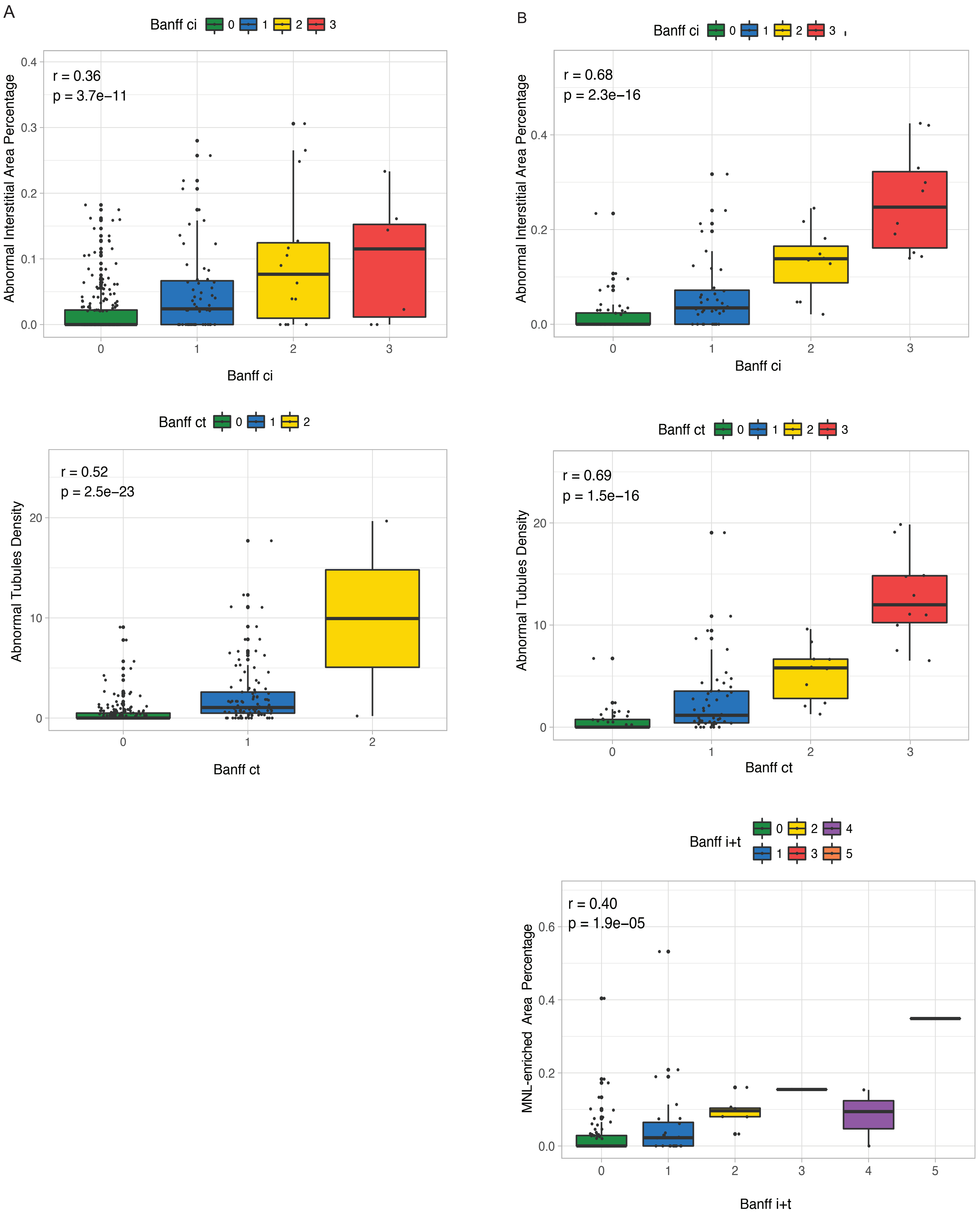

**Figure S2. Correlation of digital features with corresponding Banff scores. A)** Correlation of Abnormal Interstitial Area Percentage and Banff ci score (upper), Abnormal Tubules Density and Banff ct score (middle) in GoCAR baseline biopsy slides (n=317). **B)** Correlation of Abnormal Interstitial Area Percentage and Banff ci score (top), Abnormal Tubules Density and Banff ct score (middle), MNL-enriched Area Percentage and Banff i+t score (bottom) in AUSCAD 12m post-transplant biopsy slides (n=111).

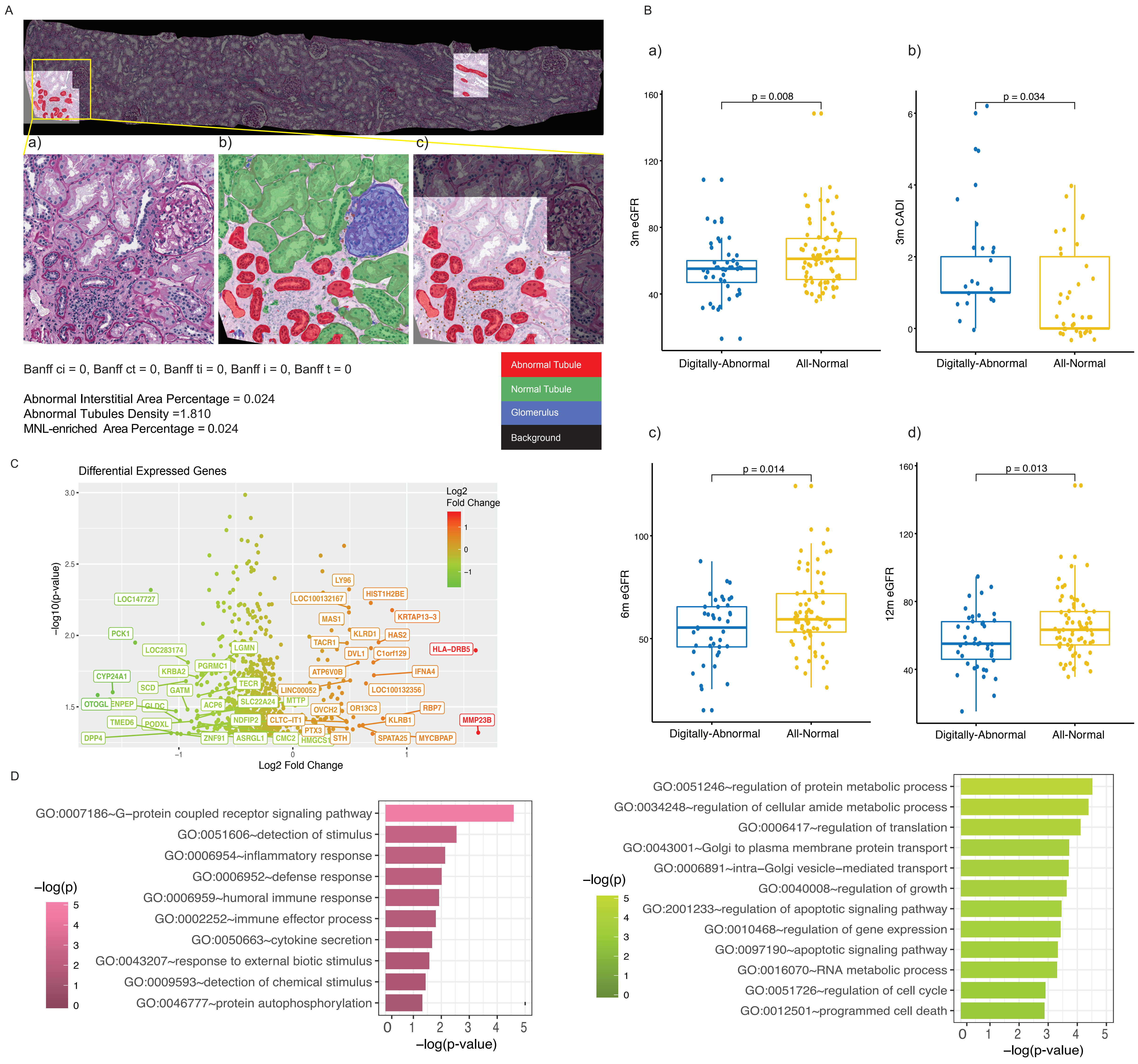

**Figure S3. Discrepancy between digital features and Banff scores. A)** An example of a WSI determined normal by all Banff scores but abnormal by digital features. Upper-panel shows whole slide image highlighting abnormal interstitium/tubules regions by digital features. Lower-panel shows close-up views of original(a), full compartment prediction(b) and abnormal ROI mask(c) from one abnormal region within yellow box. **B)** Comparison of subsequent clinical outcomes 3m eGFR(a), 3m CADI(b), 6m eGFR(c), 12m eGFR(d) between digitally-abnormal vs. –normal patients who were all determined normal by Banff scores from GoCAR baseline biopsies. P-values are calculated by t-test. **C)** Volcano plot of top 50 differentially expressed genes of GoCAR 3m post-transplant biopsy in digitally-abnormal vs.–normal patients who were all determined normal by Banff scores from GoCAR 12m biopsies. **D)** Top enriched Gene Ontology functions by up-regulated (pink, left-panel) and down-regulated (green, right-panel) genes in **(C)** comparison. P-values are calculated by Fisher's exact test.

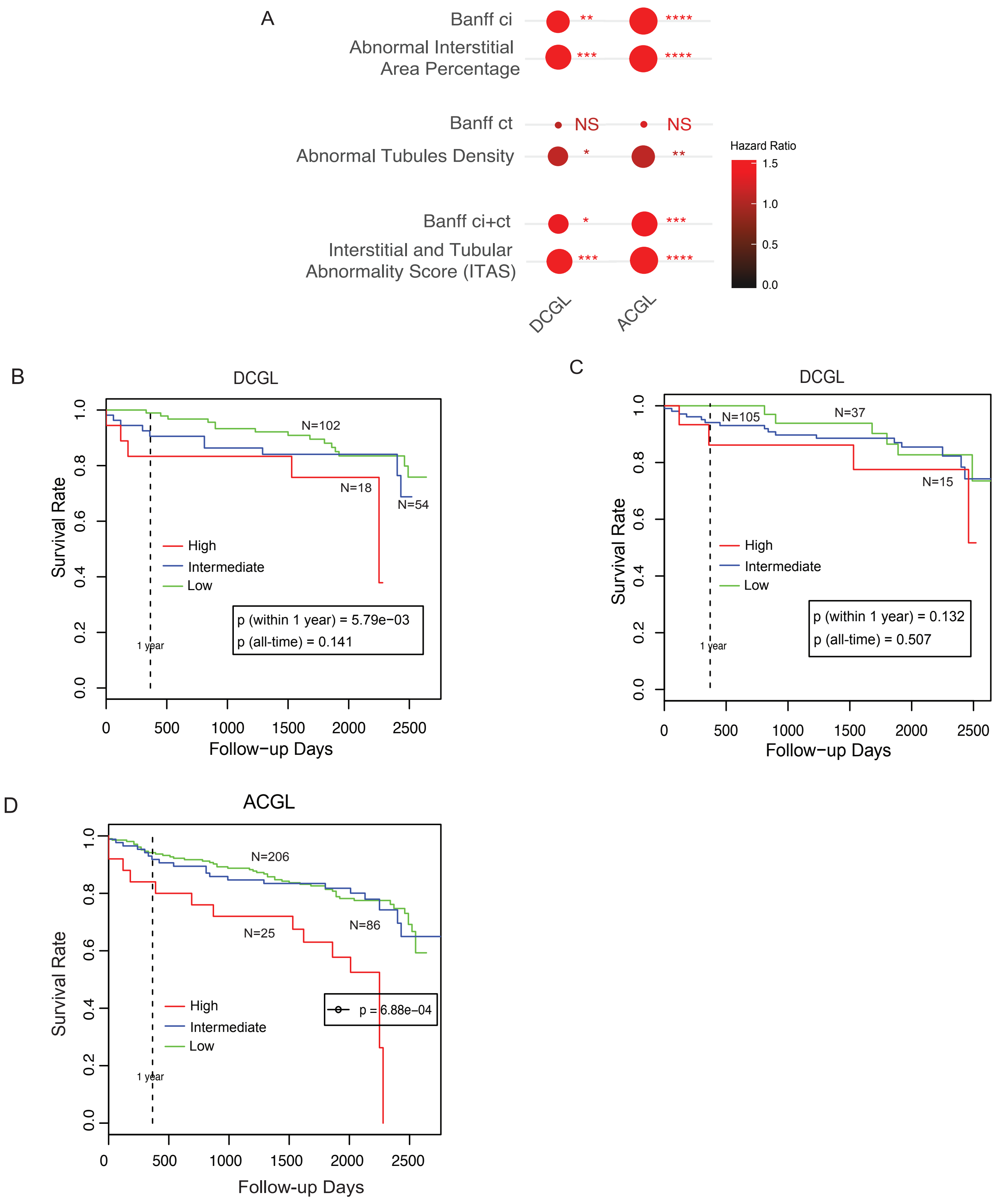

**Figure S4. Association of baseline digital features with post-transplant graft outcomes in GoCAR cohort. A)** Dot heatmap of association of Banff scores and digital features with post-transplant graft loss (death-censored graft loss (DCGL) and all-cause graft loss (ACGL)) in baseline biopsy slides (n=317). The size of dots and number of asterisks indicate significance level (p-value) of association by Cox proportional hazards regression (NS: $p \geq 0.1$; .: $0.05 \leq p < 0.1$; *:$0.005 \leq p < 0.05$; **: $5e\text{-}04 \leq p < 0.005$; ***: $5e\text{-}05 \leq p < 5e\text{-}04$; ****:$p < 5e\text{-}05$). Color darkness of dots indicate hazard ratio. **B)** Kaplan-Meier curves of DCGL in ITAS high vs. intermediate vs. low group in deceased donor population in baseline biopsies (n=174). Baseline ITAS groups are defined as: high: ITAS>0.6, intermediate: 0.1<ITAS≤0.6, low: ITAS≤0.1. P-values are calculated by log-rank test. **C)** Kaplan-Meier curves of DCGL in KDPI high vs. intermediate vs. low group in deceased donor population in baseline biopsies. KDPI groups are defined in deceased-donor population as: high: KDPI>85%, intermediate: 20%<KDPI≤85%, low: KDPI≤20%. P-values are calculated by log-rank test. **D**) Kaplan-Meier curves of ACGL in ITAS high vs. intermediate vs. low group from baseline biopsies (n=317). P-value is calculated by log-rank test.

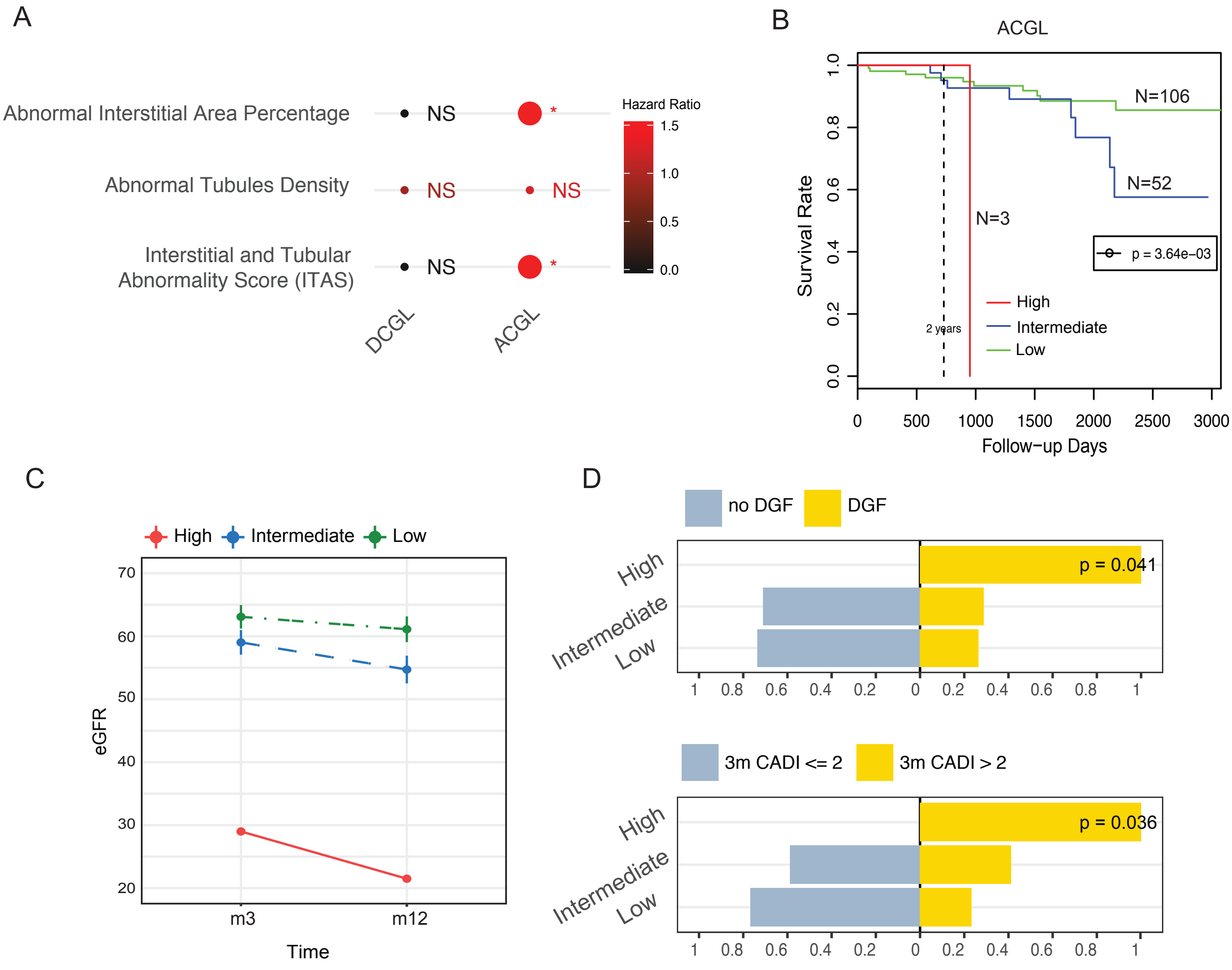

**Figure S5. Association of baseline digital features with post-transplant graft outcomes in AUSCAD cohort. A)** Dot heatmap of association of digital features with post-transplant graft loss (death-censored graft loss (DCGL) and all-cause graft loss (ACGL)) in baseline biopsy slides (n=161). The size of dots dot and number of asterisks indicate significance level (p-value) of association by Cox proportional hazards regression (NS: $p \geq 0.1$; .: $0.05 \leq p < 0.1$; *:$0.005 \leq p < 0.05$; **: 5e-04 $\leq p <$ 0.005; ***: 5e-05 $\leq p <$ 5e-04; ****:$p <$ 5e-05). Color darkness of dots indicate hazard ratio. **B)** Kaplan-Meier curves of ACGL in baseline ITAS high vs. intermediate vs. low group from baseline biopsies (n=161). Baseline ITAS groups are defined as:

high: ITAS>0.6, intermediate: 0.1<ITAS≤0.6, low: ITAS≤0.1. P-value is calculated by log-rank test. **C)** Average eGFR values over time within 12m post-transplant per baseline ITAS risk group. Error bars represent 0.1x standard deviation from mean values. **D)** Bar charts demonstrating proportions of DGF/no DGF (upper) and 3m post-transplant CADI >2/≤2 (lower) among three baseline ITAS risk groups in whole population. P-values are calculated by Fisher's exact test.

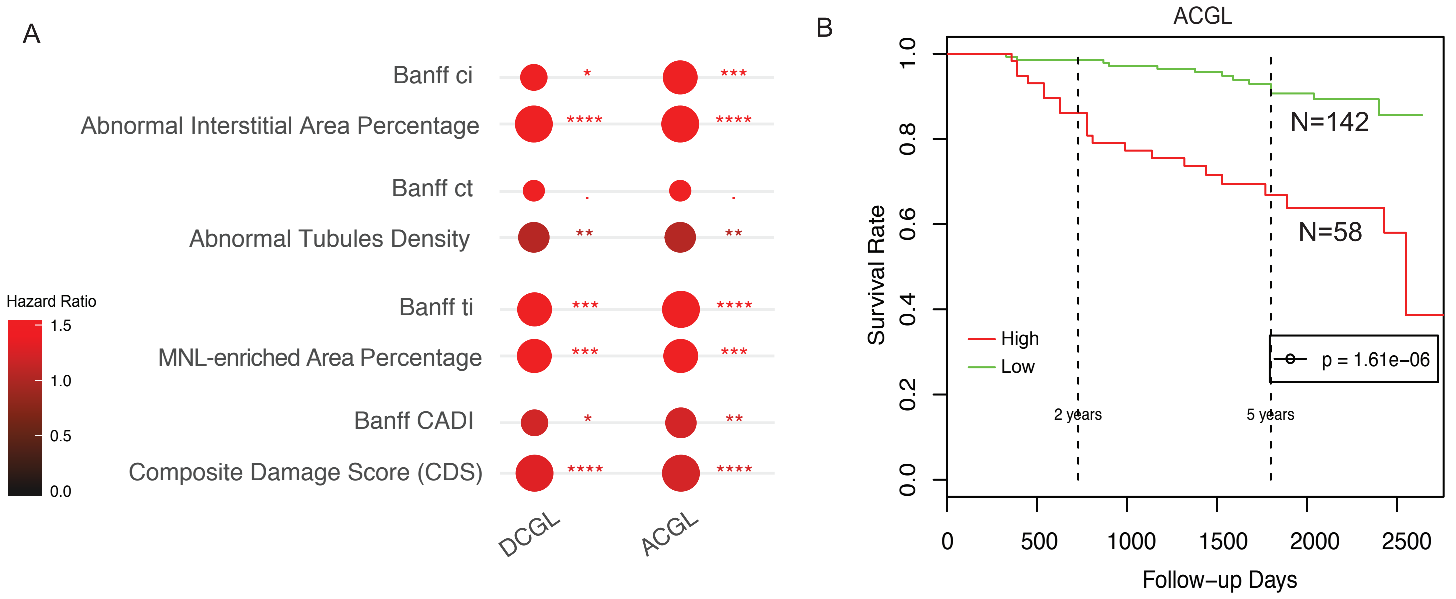

**Figure S6. Association of 12m post-transplant digital features with post-transplant graft outcomes in GoCAR cohort. A)** Dot heatmap of association of Banff scores and digital features with post-transplant graft loss (death-censored graft loss (DCGL) and all-cause graft loss (ACGL)) in 12m post-transplant biopsy slides (n=200). The size of dot and number of asterisks indicate significance level (p-value) of association by Cox proportional hazards regression (NS: $p \geq 0.1$; .: $0.05 \leq p < 0.1$; *:$0.005 \leq p < 0.05$; **: $5e\text{-}04 \leq p < 0.005$; ***: $5e\text{-}05 \leq p < 5e\text{-}04$; ****:$p < 5e\text{-}05$). Color darkness of dots indicate hazard ratio. **B**) Kaplan-Meier curves of ACGL in CDS high vs. low group from 12m post-transplant biopsies (n=200). 12m CDS groups are defined as: high: CDS>1.5, low: CDS≤1.5. P-value is calculated by log-rank test.

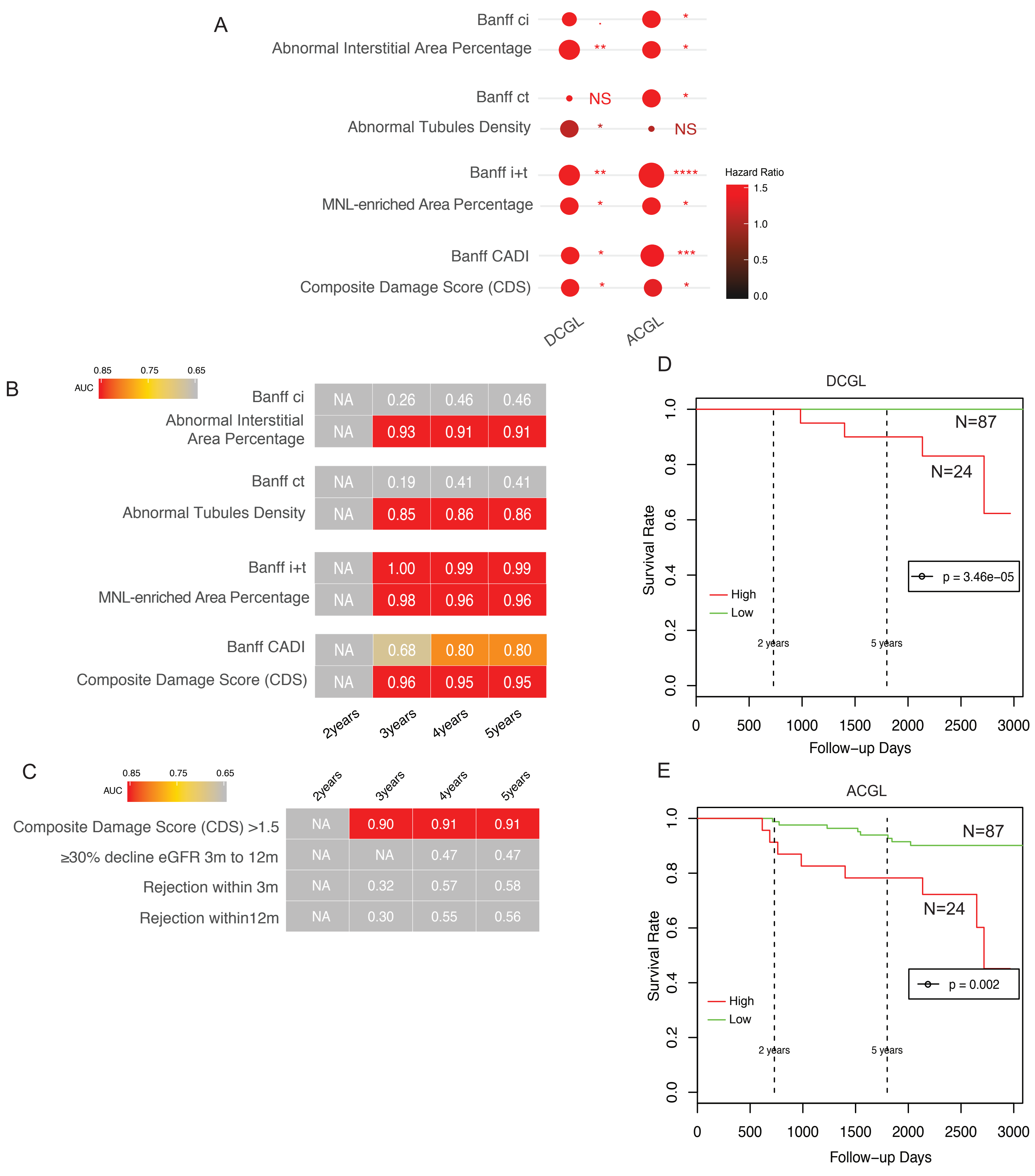

**Figure S7. Association of 12m post-transplant digital features with post-transplant graft outcomes in AUSCAD cohort. A)** Dot heatmap of association between Banff scores and digital features with post-transplant graft loss (death-censored graft loss (DCGL) and all-cause graft loss (ACGL)) in 12m post-transplant biopsy slides (n=111). The size of dot and number of asterisks indicate significance level (p-value) of association by Cox proportional hazards regression (NS: $p \geq 0.1$; .: $0.05 \leq p < 0.1$; *:$0.005 \leq p < 0.05$; **: $5e\text{-}04 \leq p < 0.005$; ***: $5e\text{-}05 \leq p < 5e\text{-}04$; ****:$p < 5e\text{-}05$). Color darkness of dots indicate hazard ratio. **B)** Heatmap of time-dependent AUCs in predicting death-censored graft loss (DCGL) by Banff scores and digital features at different time intervals in 12m post-transplant biopsy slides (n=111). Numbers and yellow-red color range of boxes represent AUC values at given time points. **C)** Heatmap of time-dependent AUCs in predicting DCGL by 12m CDS high/low group and other clinical factors which were obtained prior to or at 12m. 12m CDS groups are defined as: high: CDS>1.5, low: CDS≤1.5. **D-E)** Kaplan-Meier curves of DCGL (**D**) and ACGL (**E**) in CDS high vs. low group from AUSCAD 12m biopsies (n=111). P-values are calculated by log-rank test.

**Table S1.** Accuracy summary of kidney tissue compartment prediction model based on independent testing set.

| Group | Area Segmentation | | | Instance Detection | | |
|---|---|---|---|---|---|---|
| | TPR | PPV | F-score | TPR | PPV | F-score |
| Interstitium | 0.73 | 0.85 | 0.75 | - | - | - |
| Glomeruli | 0.94 | 0.87 | 0.93 | 0.96 | 0.97 | 0.96 |
| All Tubule | 0.93 | 0.84 | 0.91 | 0.91 | 0.85 | 0.90 |
| Normal Tubule | 0.92 | 0.79 | 0.89 | 0.81 | 0.77 | 0.80 |
| Abnormal Tubule | 0.79 | 0.78 | 0.79 | 0.84 | 0.76 | 0.82 |
| Artery | 0.84 | 0.96 | 0.86 | 0.75 | 0.89 | 0.77 |
| MN Leukocyte | - | - | - | 0.77 | 0.66 | 0.75 |
| Epithelial cell | - | - | - | 0.90 | 0.67 | 0.84 |

**Table S2. Association of baseline Banff scores and digital features with graft loss in GoCAR cohort.**

a) Association of baseline Banff scores and digital features with death-censored graft loss (DCGL) and all-cause graft loss (ACGL). P-values are calculated by Wald test from Cox proportional hazards regression.

| Scores | DCGL_pvalue | DCGL_hazard ratio | ACGL_pvalue | ACGL_hazard ratio |
|---|---|---|---|---|
| **Banff ci** | 5.0E-04 | 1.81 | 4.5E-07 | 1.82 |
| **Abnormal Interstitial Area Percentage** | 5.7E-05 | 3338.65 | 2.6E-05 | 657.78 |
| **Banff ct** | 8.1E-01 | 1.08 | 3.0E-01 | 1.26 |
| **Abnormal Tubules Density** | 1.2E-02 | 1.11 | 4.2E-03 | 1.09 |
| **Banff ci+ct** | 2.8E-02 | 1.38 | 1.1E-04 | 1.46 |
| **Interstitial and Tubular Abnormality Score (ITAS)** | 1.5E-04 | 3.25 | 3.9E-05 | 2.67 |

b) Association of baseline Banff scores and digital features with death-censored graft loss (DCGL) and all-cause graft loss (ACGL) after adjusting for clinical confounders. P-values are calculated by Wald test from Cox proportional hazards regression.

| Scores | DCGL_pvalue | DCGL_hazard ratio | ACGL_pvalue | ACGL_hazard ratio |
|---|---|---|---|---|
| **Banff ci** | 5.3E-03 | 1.62 | 4.2E-05 | 1.64 |
| **Abnormal Interstitial Area Percentage** | 1.2E-03 | 884.89 | 1.5E-03 | 160.42 |
| **Banff ct** | 6.4E-01 | 1.17 | 1.9E-01 | 1.34 |
| **Abnormal Tubules Density** | 3.1E-02 | 1.09 | 1.6E-02 | 1.08 |
| **Banff ci+ct** | 6.1E-02 | 1.31 | 7.7E-04 | 1.39 |
| **Interstitial and Tubular Abnormality Score (ITAS)** | 2.1E-03 | 2.67 | 1.4E-03 | 2.19 |

**Table S3. Association of baseline digital features with graft loss in AUSCAD cohort.**

a) Association of baseline digital features with death-censored graft loss (DCGL) and all-cause graft loss (ACGL). P-values are calculated by Wald test from Cox proportional hazards regression.

| Scores | DCGL_pvalue | DCGL_hazard ratio | ACGL_pvalue | ACGL_hazard ratio |
|---|---|---|---|---|
| **Abnormal Interstitial Area Percentage** | 3.1E-01 | 3.3E-11 | 5.0E-02 | 22674.83 |
| **Abnormal Tubules Density** | 7.6E-01 | 0.91 | 1.2E-01 | 1.23 |
| **Interstitial and Tubular Abnormality Score (ITAS)** | 3.5E-01 | 4.6E-03 | 4.6E-02 | 6.29 |

b) Association of baseline digital features with death-censored graft loss (DCGL) and all-cause graft loss (ACGL) after adjusting for clinical confounders. P-values are calculated by Wald test from Cox proportional hazards regression.

| Scores | DCGL_pvalue | DCGL_hazard ratio | ACGL_pvalue | ACGL_hazard ratio |
|---|---|---|---|---|
| **Abnormal Interstitial Area Percentage** | 2.0E-01 | 2.7E-14 | 4.7E-02 | 38710.63 |
| **Abnormal Tubules Density** | 5.0E-01 | 0.79 | 1.8E-01 | 1.20 |
| **Interstitial and Tubular Abnormality Score (ITAS)** | 2.4E-01 | 7.5E-04 | 5.2E-02 | 6.17 |

**Table S4. Association of 12m post-transplant Banff scores and digital features with graft loss in GoCAR cohort.**

a) Association of 12m Banff scores and digital features with death-censored graft loss (DCGL) and all-cause graft loss (ACGL). P-values are calculated by Wald test from Cox proportional hazards regression.

| Scores | DCGL_pvalue | DCGL_hazard ratio | ACGL_pvalue | ACGL_hazard ratio |
|---|---|---|---|---|
| **Banff ci** | 1.6E-02 | 1.74 | 2.9E-04 | 1.86 |
| **Abnormal Interstitial Area Percentage** | 2.1E-05 | 147.43 | 1.2E-05 | 67.65 |
| **Banff ct** | 9.6E-02 | 1.62 | 5.0E-02 | 1.56 |
| **Abnormal Tubules Density** | 1.5E-03 | 1.06 | 1.4E-03 | 1.05 |
| **Banff ti** | 2.2E-04 | 2.10 | 3.9E-05 | 1.89 |
| **MNL-enriched Area Percentage** | 1.6E-04 | 25.48 | 2.9E-04 | 13.09 |
| **Banff CADI** | 2.2E-02 | 1.22 | 2.2E-03 | 1.23 |
| **Composite Damage Score (CDS)** | 1.8E-05 | 1.30 | 2.5E-05 | 1.24 |

b) Association of 12m Banff scores and digital features with death-censored graft loss (DCGL) and all-cause graft loss (ACGL) after adjusting for clinical confounders. P-values are calculated by Wald test from Cox proportional hazards regression.

| Scores | DCGL_pvalue | DCGL_hazard ratio | ACGL_pvalue | ACGL_hazard ratio |
|---|---|---|---|---|
| **Banff ci** | 5.2E-02 | 1.62 | 2.0E-03 | 1.78 |
| **Abnormal Interstitial Area Percentage** | 2.7E-04 | 101.95 | 1.9E-04 | 44.57 |
| **Banff ct** | 2.3E-01 | 1.44 | 1.4E-01 | 1.42 |
| **Abnormal Tubules Density** | 9.5E-03 | 1.05 | 1.0E-02 | 1.04 |
| **Banff ti** | 3.5E-03 | 1.86 | 1.5E-03 | 1.67 |
| **MNL-enriched Area Percentage** | 3.7E-03 | 13.04 | 9.1E-03 | 6.65 |
| **Banff CADI** | 1.4E-01 | 1.15 | 3.9E-02 | 1.16 |
| **Composite Damage Score (CDS)** | 7.2E-04 | 1.25 | 1.4E-03 | 1.18 |

**Table S5. Association of 12m post-transplant Banff scores and digital features with graft loss in AUSCAD cohort.**

a) Association of 12m Banff scores and digital features with death-censored graft loss (DCGL) and all-cause graft loss (ACGL). P-values are calculated by Wald test from Cox proportional hazards regression.

| Scores | DCGL_pvalue | DCGL_hazard ratio | ACGL_pvalue | ACGL_hazard ratio |
|---|---|---|---|---|
| **Banff ci** | 9.0E-02 | 2.20 | 1.5E-02 | 1.72 |
| **Abnormal Interstitial Area Percentage** | 4.9E-03 | 2461.82 | 9.5E-03 | 52.17 |
| **Banff ct** | 1.1E-01 | 2.17 | 1.7E-02 | 1.76 |
| **Abnormal Tubules Density** | 4.2E-02 | 1.10 | 1.0E-01 | 1.05 |
| **Banff i+t** | 2.2E-03 | 6.32 | 3.8E-07 | 2.26 |
| **MNL-enriched Area Percentage** | 8.8E-03 | 2019.32 | 7.3E-03 | 93.71 |
| **Banff CADI** | 5.6E-03 | 1.91 | 2.1E-04 | 1.41 |
| **Composite Damage Score (CDS)** | 6.1E-03 | 1.71 | 6.5E-03 | 1.33 |

b) Association of 12m Banff scores and digital features with death-censored graft loss (DCGL) and all-cause graft loss (ACGL) after adjusting for clinical confounders. P-values are calculated by Wald test from Cox proportional hazards regression.

| Scores | DCGL_pvalue | DCGL_hazard ratio | ACGL_pvalue | ACGL_hazard ratio |
|---|---|---|---|---|
| **Banff ci** | 8.1E-02 | 2.30 | 5.6E-03 | 1.92 |
| **Abnormal Interstitial Area Percentage** | 7.0E-03 | 2220.88 | 9.3E-03 | 59.92 |
| **Banff ct** | 8.7E-02 | 2.32 | 3.5E-03 | 2.02 |
| **Abnormal Tubules Density** | 7.3E-02 | 1.09 | 1.5E-01 | 1.05 |
| **Banff i+t** | 2.0E-02 | 11.63 | 1.1E-06 | 2.46 |
| **MNL-enriched Area Percentage** | 2.1E-02 | 1165.56 | 1.4E-02 | 66.25 |
| **Banff CADI** | 5.1E-03 | 1.94 | 2.2E-04 | 1.46 |
| **Composite Damage Score (CDS)** | 1.4E-02 | 1.67 | 1.2E-02 | 1.32 |

1. O'Connell, P.J., et al., *Biopsy transcriptome expression profiling to identify kidney transplants at risk of chronic injury: a multicentre, prospective study.* Lancet, 2016. **388**(10048): p. 983-93.
2. He, K., et al. *Mask R-CNN*. 2017. arXiv:1703.06870.
3. Ronneberger, O., P. Fischer, and T. Brox *U-Net: Convolutional Networks for Biomedical Image Segmentation*. 2015. arXiv:1505.04597.
4. Abdulla, W. *Mask R-CNN for object detection and instance segmentation on Keras and TensorFlow*. GitHub repository 2017; Available from: https://github.com/matterport/Mask_RCNN.
5. Akeret, J., et al., *Radio frequency interference mitigation using deep convolutional neural networks.* Astronomy and Computing, 2017. **18**: p. 35.
6. Van Rijsbergen, C.J., *Information Retrieval (2nd ed.).* . 1979: Butterworth-Heinemann.
7. Ritchie, M.E., et al., *limma powers differential expression analyses for RNA-sequencing and microarray studies.* Nucleic Acids Research, 2015. **43**(7).
8. Huang da, W., B.T. Sherman, and R.A. Lempicki, *Systematic and integrative analysis of large gene lists using DAVID bioinformatics resources.* Nat Protoc, 2009. **4**(1): p. 44-57.
